# Spatial suppression in visual motion perception is driven by inhibition: evidence from MEG gamma oscillations

**DOI:** 10.1101/861765

**Authors:** E.V. Orekhova, E.N. Rostovtseva, V.O. Manyukhina, A.O. Prokofiev, T.S. Obukhova, A.Yu. Nikolaeva, J.F. Schneiderman, T.A. Stroganova

## Abstract

Spatial suppression (SS) is a visual perceptual phenomenon that is manifest in a reduction of directional sensitivity for drifting high-contrast gratings whose size exceeds the center of the visual field. Gratings moving at faster velocities induce stronger SS. The neural processes that give rise to such size- and velocity-dependent reductions in directional sensitivity are currently unknown, and the role of surround inhibition is unclear. In magnetoencephalogram (MEG), large high-contrast drifting gratings induce a strong gamma response (GR), which also attenuates with an increase in the gratings’ velocity. It has been suggested that the slope of this GR attenuation is mediated by inhibitory interactions in the primary visual cortex. Herein, we investigate whether SS is related to this inhibitory-based MEG measure. We evaluated SS and GR in two independent samples of participants: school-age boys and adult women. The slope of GR attenuation predicted inter-individual differences in SS in both samples. Test-retest reliability of the neuro-behavioral correlation was assessed in the adults, and was high between two sessions separated by several days or weeks. Neither frequencies nor absolute amplitudes of the GRs correlated with SS, which highlights the functional relevance of velocity-related *changes* in GR magnitude caused by augmentation of incoming input. Our findings provide evidence that links the psychophysical phenomenon of SS to inhibitory-based neural responses in the human primary visual cortex. This supports the role of inhibitory interactions as an important underlying mechanism for spatial suppression.

**Highlights:**

- The role of surround inhibition in perceptual spatial suppression (SS) is debated
- GR attenuation with increasing grating’s velocity may reflect surround inhibition
- People with greater GR attenuation exhibit stronger SS
- The neuro-behavioral correlation is replicated in school-age boys and adult women
- The surround inhibition in the V1 is an important mechanism underlying SS

## 1. Introduction

Vision is the main source of sensory information in humans. Considering the abundance of visual information bombarding the brain at any given moment, visual processing is inherently competitive and requires neural mechanisms that suppress redundant or distracting input (Desimone and Duncan, 1995). It has been suggested that, for efficient information processing, the brain applies a so called ‘sparse coding’ strategy, i.e. encoding of sensory information using small number of active neurons (Baddeley et al., 1997; Barlow, 1961; Field, 1987; Olshausen and Field, 2004). Surround suppression – weakening of neurons’ responses to the combined stimulation of their classical receptive fields and surrounding periphery – is an important mechanism of sparse coding (Angelucci et al., 2017; Angelucci and Bressloff, 2006; Sachdev et al., 2012; Zhu and Rozell, 2013). In the visual cortex, surround suppression improves coding efficiency of natural images by promoting figure-ground segmentation of moving objects (Tadin et al., 2019) and by facilitating the processing of potentially salient peripheral motion (Nurminen and Angelucci, 2014).

These perceptual advantages, however, come at a price, because surround suppression is thought to lead to diminished directional sensitivity to drifting high-contrast regular shapes, such as gratings, if these shapes extend beyond the center of the visual field (Tadin, 2015; Tadin et al., 2003). This visual perceptual phenomenon, called *spatial suppression* (SS), has been found in both humans (Tadin, 2015; Tadin et al., 2003) and nonhuman primates (Liu et al., 2016). In psychophysical experiments, SS manifests itself in a longer time needed to discriminate the direction of motion of large high-contrast gratings, compared to small ones. Reduced SS has been posited as a behavioral indication of impaired neural surround inhibition in old age (Betts et al., 2007, 2012; Tadin et al., 2019) and in a number of neuropsychiatric disorders, such as depression (Golomb et al., 2009), schizophrenia (Tadin et al., 2006) and ASD (Foss-Feig et al., 2013; Sysoeva et al., 2017). However, the role of neural surround inhibition in SS in humans remains elusive and is a subject of continuing debate (see e.g. Schallmo et al., 2018).

The role of early and higher-tier visual cortical areas in the psychophysical phenomenon of SS is also unclear. SS has often been attributed to neuronal surround suppression within area MT (Tadin, 2015; Tadin et al., 2011), which plays a major role in motion direction sensitivity (Born and Bradley, 2005). Tadin and colleagues have shown that TMS-induced periods of reduced excitability in MT were associated with weakening of SS, and suggested that such disruptive TMS is interfering with inhibitory processing within MT/V5, which, in turn, improves perception of large moving stimuli (Tadin et al., 2011). Contrasting this view, studies in monkeys demonstrate that the strength of neuronal surround suppression in the MT is low and cannot explain the reported high level of deterioration in behavioral performance (Liu et al., 2016).

A recent human study combining behavioral psychophysics, functional MRI and magnetic resonance spectroscopy (Schallmo et al., 2018) further challenged the proposed role of heightened inhibitory transmission within MT in SS. Although increasing the size of moving gratings did suppress fMRI responses in MT, the suppression was much more subtle than, and could thus be considered secondary to, that in the primary visual cortex (V1). Further, the authors did not find the expected direct association between SS measured in a psychophysical task and the concentration of the inhibitory neurotransmitter GABA in either area MT or the early visual cortex. In a similar vein, GABA concentration was unrelated to the fMRI response suppression caused by increasing the size of drifting gratings. Therefore, Schallmo et al. came to the conclusion that SS is not primarily driven by GABA-mediated inhibition.

Given the complex dynamic nature of the excitatory-inhibitory interactions elicited by large moving gratings that has been revealed in animal studies (Angelucci et al., 2017; Angelucci and Bressloff, 2006; Nurminen et al., 2018), individual variations in the baseline ‘bulk’ GABA concentration in visual areas might be only one of many factors contributing to SS. The better way to clarify the role of GABAergic inhibition in human SS could be to utilize a measure of how efficiently the *‘on-demand’* release of inhibition in V1 follows the increase in excitatory input. This can be achieved through recording magnetoencephalographic (MEG) visually-induced gamma oscillations that reflect the fast collective fluctuations in membrane potentials of V1 neurons (Buzsaki et al., 2012).

Cortical inhibition is essential for generation of neural oscillations in the gamma frequency range - 30-100 Hz (Atallah et al., 2012; Buzsaki and Wang, 2012; Cardin et al., 2009; Hasenstaub et al., 2005; Sohal et al., 2009; Takada et al., 2014). Recent findings by our group suggest that the efficiency of inhibitory control in the human V1 is reflected in degree of *attenuation* of the MEG gamma response (GR) with increasing velocity of large high-contrast visual gratings (Orekhova et al., 2019; Orekhova et al., 2018b). This GR attenuation was accompanied by an increase in gamma frequency, which presumably reflects an increase in tonic excitation of V1 inhibitory neurons (Anver et al., 2011; Mann and Mody, 2010). An important factor leading to velocity-related attenuation of the MEG GR seems to be changes in bottom-up and/or top-down excitatory drive to V1 inhibitory cells. Indeed, it has been shown that the ‘too strong’ drive disrupts gamma synchrony (Borgers and Kopell, 2005; Cannon et al., 2014; King et al., 2013; Kopell et al., 2010). On the other hand, the resulting asynchronous inhibition especially effectively weakens the activation of excitatory cells in response to high excitatory input (Borgers and Kopell, 2005). Therefore, at the individual level, the degree of GR attenuation may reflect how efficiently neural inhibition mitigates intensive excitatory input to V1 and reduces the network excitatory response.

In the present MEG study, we tested for the link between the degree of motion-related attenuation of the visual GR and SS in human visual motion perception. We hypothesized that if both GR attenuation and SS reflect the efficiency of surround inhibition in the human visual cortex, then the interindividual variation in the GR attenuation should mirror changes in directional sensitivity observed with increasing the size of drifting gratings. Therefore, we predicted that those individuals who have stronger GR attenuation (i.e., highly efficient down-regulation of V1 excitation) would exhibit worse directional sensitivity *for the large, but not for the small*, moving gratings. This precise prediction allows us to rule out the other neural factors, e.g., attention focusing and general motion sensitivity that, unlike surround inhibition, should have a similar impact on the correlations between GR and psychophysical thresholds for both small and large gratings. The presence of such a selective correlation would therefore provide strong support for a causal link between excitatory/inhibitory balance in V1 and the perceptual phenomenon of SS.

It has been widely discussed that cognitive neuroscience suffers from a ‘replication crisis’ due to small sample sizes and a bias to publish positive findings (Button et al., 2013; Szucs and Ioannidis, 2017). To decrease the probability of ‘false-positive’ results in our study, we tested for the predicted neuro-behavioral correlation in the two diverse samples of individuals. In addition to previously collected data from school-age children (all boys) (Stroganova et al., 2018), we performed the same MEG and psychophysical measurements in a sample of adult women. Given the fundamental nature of the proposed inhibitory-based mechanism, it should influence both perception and gamma oscillatory dynamics in adults and children irrespective of gender. We reasoned that apart from testing for reproducibility, replication of the anticipated neuro-behavioral relationship in these demographically diverse samples would provide a rigorous test of its functional relevance. To furthermore ensure temporal stability of the target measures, we also estimated their test/re-test reliability by repeating the measurements on the adult individuals in two sessions that were separated by at several days or weeks.

## 2. Materials and Methods

### 2.1. Participants

*Sample 1* included 18 healthy women (aged 18-40 years, mean 28 years) who were a part of another study. All the participants underwent the psychophysical testing and MEG recording procedure twice with an interval of 5-118 days (at the follicular and luteal stages of the menstrual cycle, the stages being counterbalanced between the visits). For the purpose of the present study, the data obtained during the 1^st^ and the 2^nd^ visits were averaged except when evaluating test-retest reliability.

*Sample 2* initially included 43 typically developing children (all boys aged 6-15 years) who participated in two other studies (Orekhova et al., 2018b; Sysoeva et al., 2017). In nine of these children, no reliable MEG GRs were detected according to the acceptance criteria (see below), which prevented reliable estimation of the gamma suppression slope. These children were excluded from the analysis. The final sample therefore comprised 34 participants (age 7.7-15.5 years, mean=11.4 years). In all of these participants, IQ was assessed using the Kaufman Assessment Battery for Children К-АВС II (Kaufman and Kaufman, 2004).

### 2.2. Psychophysical task

To estimate SS, we used a psychophysical task that has been previously applied in healthy individuals (Arranz-Paraiso and Serrano-Pedraza, 2018; Betts et al., 2012; Melnick et al., 2013; Tadin et al., 2003; Troche et al., 2018) as well as in those with neurological or psychiatric disorders (Foss-Feig et al., 2013; Golomb et al., 2009; Sysoeva et al., 2017; Tadin et al., 2006). A detailed description of the current version of the experimental procedure is given in the (Sysoeva et al., 2017).

Visual stimuli were presented using PsychToolbox (Brainard, 1997). Figure 1 illustrates the types of experimental stimuli and the procedure we used. The stimuli consisted of a vertical full-contrast sine wave grating (1 cycle/°) that drifted at a constant speed of 4 °/s. The size of the stimuli was controlled by a two-dimensional Gaussian envelope whose full-width at half-maximum was set to 1, 2.5, or 12° for the small, medium, and large stimuli types. Participants were instructed to discriminate the direction of motion. An automated staircase procedure was used in order to estimate the minimal exposure time needed for correct motion direction discrimination. In adults, we used three types of stimuli (large: 12° of visual angle, medium: 2.5°, and small: 1° gratings), while only the large and the small gratings were presented to the children. The direction of visual motion (left or right) was determined randomly for each trial. Participants’ eyes were ~60 cm from the monitor on which stimuli were presented (Benq XL2420T, 24′′W LED, 1,920 Å~ 1,080 resolution, 120 Hz).

**Fig. 1.**
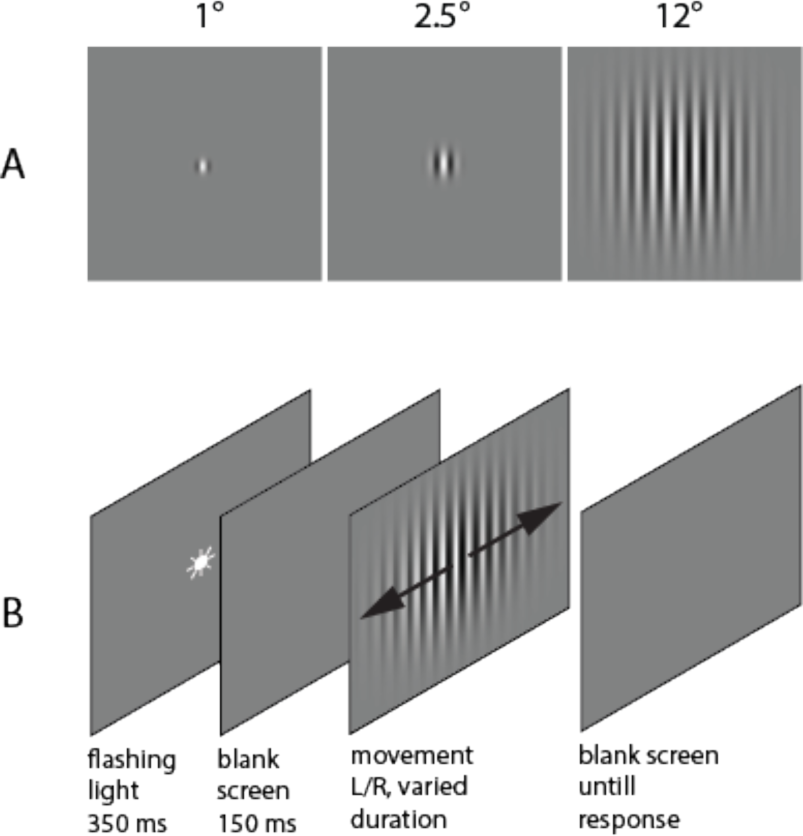
Psychophysical experiment: stimuli (A) and schematic representation of the experimental paradigm (B).

To estimate the strength of SS, we calculated the spatial suppression index (SSI) via: SSI= log10[Threshold_LARGE_/Threshold_SMALL_]

The experimental procedure was explained in detail to each participant. Children also performed a short training session before the main experiment. During the experiment, a research assistant that was seated next to the child maintained the correct distance from the monitor, vertical head position, and adequate task performance. Participants were asked to make an un-speeded two-alternative forced-choice response that indicated the perceived right or left direction of motion by pressing the left or right arrows on a keyboard, respectively. The inter-trial interval was 500 ms. In the beginning of each trial, a central dot flickered on the screen (50 ms on, 50 ms off, 250 ms on, 150 ms off) followed by the stimulus presentation.

In children the initial stimulus duration was set to 150 ms and the duration was changed with a fixed step size of 8.3 ms. The duration was further adjusted depending on the participant’s response, using an interleaved one-up two down staircase approach that aimed to converge on a 71% correct performance. Independent staircases were used for each type of stimuli. The block continued until all staircases completed 9 reversals. The thresholds were computed by averaging over the last 7 reversals. The participants repeated the block two times in a row and the thresholds obtained in the two blocks were averaged. The procedure applied for adults was identical to that for children with minor exceptions. First, the initial stimulus duration in adults was set at 200 ms. Second, the initial step in adults was set at 16.7 ms and decreased to 8.3 ms after the first two reversals.

### 2.3. MEG Experiment

#### Experimental task

We applied the experimental paradigm that has been shown to induce reliable MEG visual GRs in previous studies (Orekhova et al., 2015; Orekhova et al., 2019; Orekhova et al., 2018b). The stimuli were generated using Presentation software (Neurobehavioral Systems Inc., The United States) and presented using a PT-D7700E-K DLP projector with 1280 х 1024 screen resolution and 60 Hz refresh rate. They consisted of black and white sinusoidally modulated annular gratings with a spatial frequency of 1.66 cycles per degree of visual angle and an outer size of 18 degrees of visual angle (Fig. 2A). The gratings appeared in the center of the screen over a black background and either remained static or drifted to the central point at velocities of 1.2, 3.6, or 6.0°/s, (which approximately corresponds to temporal frequencies of 0, 2, 6, and 10 Hz, respectively); hereafter, we respectively refer to these stimuli as ‘static’, ‘slow’, ‘medium,’ and ‘fast.’ In children, only the moving stimuli were presented.

**Fig. 2.**
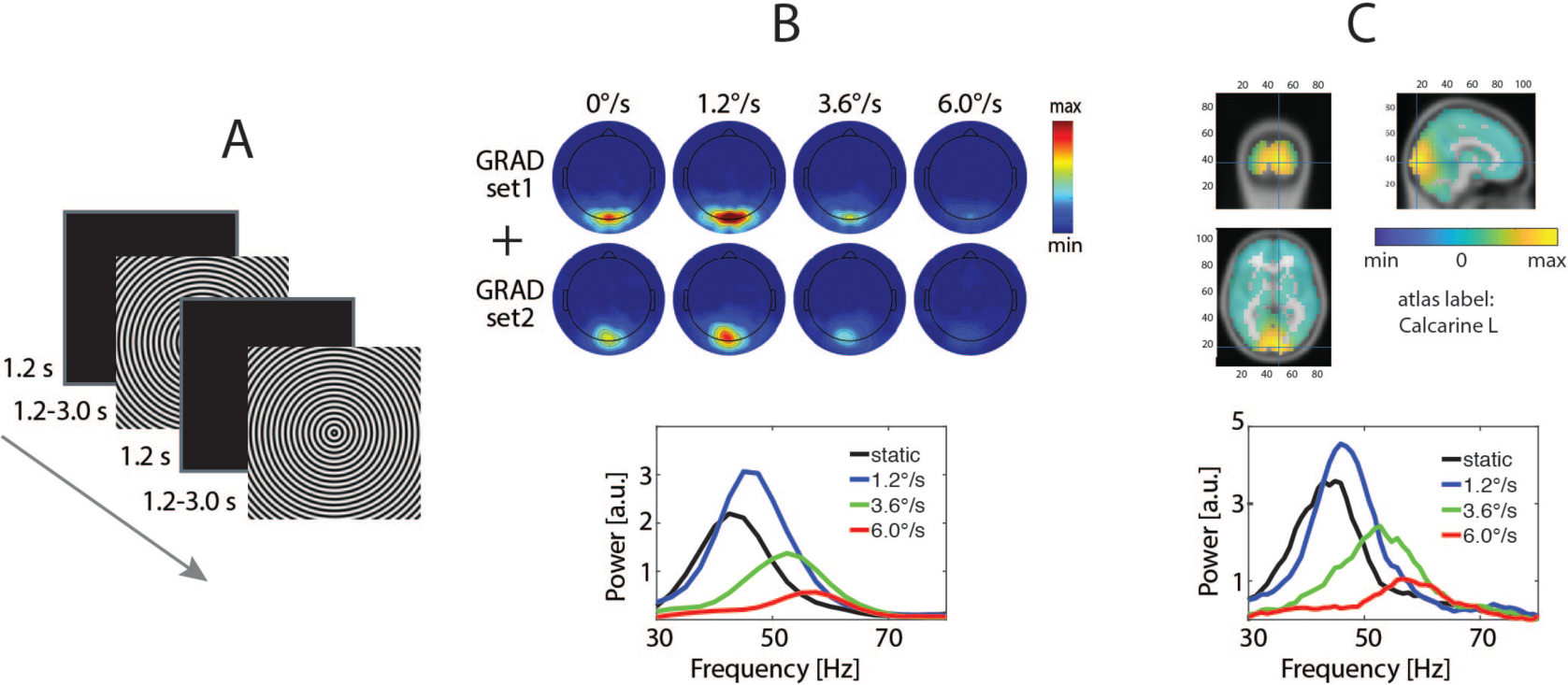
MEG experiment: stimuli and data processing. **A.** Experimental paradigm. The gratings were shown for 1.2-3 sec while they either remained static or drifted to the center of the screen immediately upon display at 1.2, 3.6, or 6.0 °/s. After the subject’s, a fixation cross appeared in the center of the black screen for 1.2 sec response (see Materials and Methods for details). **B.** GR analysis in sensor space: results for a representative subject. The average change in spectral power (relative to baseline) was calculated from the gradiometer pair that showed the maximal broadband stimulus-related gamma increase. **C.** GR in the source space: results for a representative subject. LCMV beamformers were used to reconstruct the signal at ‘virtual sensors’ inside the brain. The average change in spectral power was calculated for the set of 26 connected vertices that included and surrounded the vertex with the maximal broadband GR (see Materials and Methods for details on vertex selection).

Each trial began with a presentation of a fixation cross in the center of the display over a black background for 1200 ms that was followed by the grating that drifted for 1200-3000 ms and then stopped. The participants were instructed to respond with a button press to changes in the stimulation stream (termination of motion for moving stimuli or disappearance of the grating for static stimuli). If no response occurred within 1 s, the grating was substituted by a discouraging message ‘too late!’ that remained on the screen for 2000 ms, after which a new trial began. Stimuli were presented in three experimental blocks in a random order, resulting in 90 repetitions of each stimulus type. The luminance of the screen measured at the position of the observer’s eyes was 53 Lux during the stimulation and 2.5 Lux during the inter-stimulus interval. Short (3–6 s) animated cartoon characters were presented randomly between every 2–5 stimuli to increase vigilance and minimize fatigue.

#### MEG recording

MEG was recorded at the Moscow MEG Centre (http://megmoscow.com/) using a 306-channel system (ElektaNeuromag Vectorview). The data was recorded with high- and low-pass filters at 0.1 and 330 Hz, respectively, and digitized at 1000 Hz. The subjects’ head position during MEG recording was continuously monitored.

#### MEG data preprocessing

The data was de-noised using the Temporal Signal-Space Separation (tSSS) method (Taulu and Hari, 2009), motion corrected, and adjusted to a common head position. For the following pre-processing, MNE-python software was used (Gramfort et al., 2013). The de-noised data was low-passed at 200 Hz and resampled at 500 Hz. Independent component analysis (ICA) was used to correct for biological artifacts. The data was then epoched (−1 to 1.2 s relative to the stimulus onset) and checked for the presence of residual artifacts. After rejection of the artifact-contaminated epochs and error trials, the average number of ‘good’ epochs per subject in children was 71, 72 and 71 for the ‘slow’, ‘medium,’ and ‘fast’ conditions. In adults the average number of good trials per subject over the two sessions was 80, 81, 79 and 80 for the ‘static’, ‘slow’, ‘medium’, and ‘fast’ conditions. More details about MEG data preprocessing are given in (Orekhova et al., 2019).

#### MEG data analysis at the sensor level

Sensor-level analysis was performed using MNE-python software. To decrease the contribution of phase-locked activity (related to the appearance of the stimulus on the screen, as well as of the effect of photic driving that could be induced by the drift rate, screen refresh rate, or by their interaction), we subtracted averaged evoked responses from each single data epoch using the ‘subtract_evoked’ MNE function.

To estimate MEG signal power, we applied multitaper analysis (tfr_multitaper) with 3 tapers (time_bandwidth = 4), 2.5 Hz frequency resolution, and the number of cycles equal to frequency/2.5. We used a fixed multitaper window of 400 ms that was centered at −800:50:-200 ms time points for the pre-stimulus interval and at 400:50:1000 ms points for the post-stimulus interval. The power of the GR was then estimated as the normalized power difference between the stimulation period (stim) and the baseline period (base): [stim-base]/base.

To locate the ‘maximal response sensor pair’, the normalized power changes were averaged over the two gradiometers of each triplet of sensors in the gamma range (50-110 Hz in children and 35-110 Hz in adults). The location of the maximal increase of the broad-band gamma power was defined among the selected posterior positions (’MEG1932/3’, ‘MEG1922/3’, ‘MEG2042/3’, ‘MEG2032/3’, ‘MEG2112/3’, ‘MEG2122/3’, ‘MEG2342/3’ and ‘MEG2332/3’), separately for each condition. The power response spectra were averaged over the two gradiometers and smoothed in frequency with a 3-point-averaging window. The peak gamma frequency and power were then defined at the ‘maximal pair’ of the gradiometers. A frequency range of interest was defined at those frequencies of the gamma range where the [stim-base]/base ratio exceeded 2/3 of the maximum for the particular subject and condition. The gamma peak frequency was calculated as the center of gravity, whereas the GR power was calculated as the average power over that range.

For each subject/condition, we calculated probabilities of a post-stimulus increase in gamma power in the stimulation period relative to the baseline using the Wilcoxon test. We considered the response reliable if the probability of the gamma power increase at its absolute maximum was significant at p < .0001.

In order to quantify the degree of suppression of the GR power with increasing visual motion velocity, we used the ‘gamma suppression slope’ (GSS) index introduced in our previous study (Orekhova et al., 2019). GSS is the coefficient of regression of the weighted GR power to velocity. To calculate GSS, we used the ‘fitlm’ Matlab function: fitlm (x, y, ‘y ~ x1–1’); where x = [1.2, 3.6, 6.0], y = [0, POWmedium/POWslow–1, POWfast/POWslow–1] and ‘y ~ x1–1’ sets the intercept of the regression line to zero. The resulting regression coefficient b is equal to zero in the case of a constant response power in the three experimental conditions (i.e., ‘no suppression’) and is proportionally more negative in case of stronger velocity-related suppression of the GR.

#### MEG data analysis at the source level

The gamma activity source location was estimated with the LCMV beamformer analysis implemented in the FieldTrip M/EEG toolbox (Oostenveld et al., 2011). Prior to analysis, each subject’s brain was morphed to the MNI template brain using a nonlinear normalization and 0.6 mm grid. The signal was filtered in the 30-120 Hz range and the ‘virtual sensors’ time series were extracted for each of 6855 brain voxels. The time-frequency analysis (‘mtmfft’ method, ±5 Hz smoothing, ~1Hz frequency resolution) was performed for pre-stimulus (−0.8 to 0 s) and stimulus (0.4 to 1.2 s) windows for each virtual source and the results were averaged over trials. The GR parameters were then computed for its spatial maximum, which was defined as the voxel with the most significant increase in 45-90 Hz power during visual stimulation and twenty five surrounding neighboring voxels (i.e. 26 voxels). To assess the source-derived GR frequency and power, we further applied the same approach as described for the sensor space analysis.

#### Statistical analysis

To assess the test-retest reliability of MEG and psychophysical parameters, we calculated intra-class correlations using SPSS (two-way mixed model, absolute-agreement type). To estimate the effect of stimulus type, the Friedman ANOVA was used. We used Spearman correlation coefficients to quantify the correlations between variables. The data tables used for statistical analysis are available as Supplementary information to this article (Supplementary Data 1-3).

## 3. Results

### 3.1. Psychophysics

#### Spatial suppression effect in adults

A Friedman ANOVA with factor Size (large, medium and small) revealed a highly reliable effect of grating size (Chi-Sqr_(18, 2)_=30.3, p<0.0001, Kendel’s W=0.84) that was due to increase of the motion direction discrimination threshold from the small to the medium and further to the large grating (Fig. 3). The effect of SS in our experimental setup was large: at the group level, we observed a more than 300% rise in the detection time as the stimulus size increased from 1 to 12° of visual angle. The median thresholds were 35 ms for the small and 117 ms for the large gratings. This was an important finding, since previous studies suggest that the SS effect may be confined to a specific range of stimulus parameters and/or certain types of displays (Troche et al., 2018). We also observed high inter-individual variability in SS: SSI values ranged from 0.05 to 1.78, i.e. a 23% to 5808% increase in duration threshold. One of the participants demonstrated extremely strong SS: she needed more than one second to identify the direction of motion for the large grating.

**Fig. 3.**
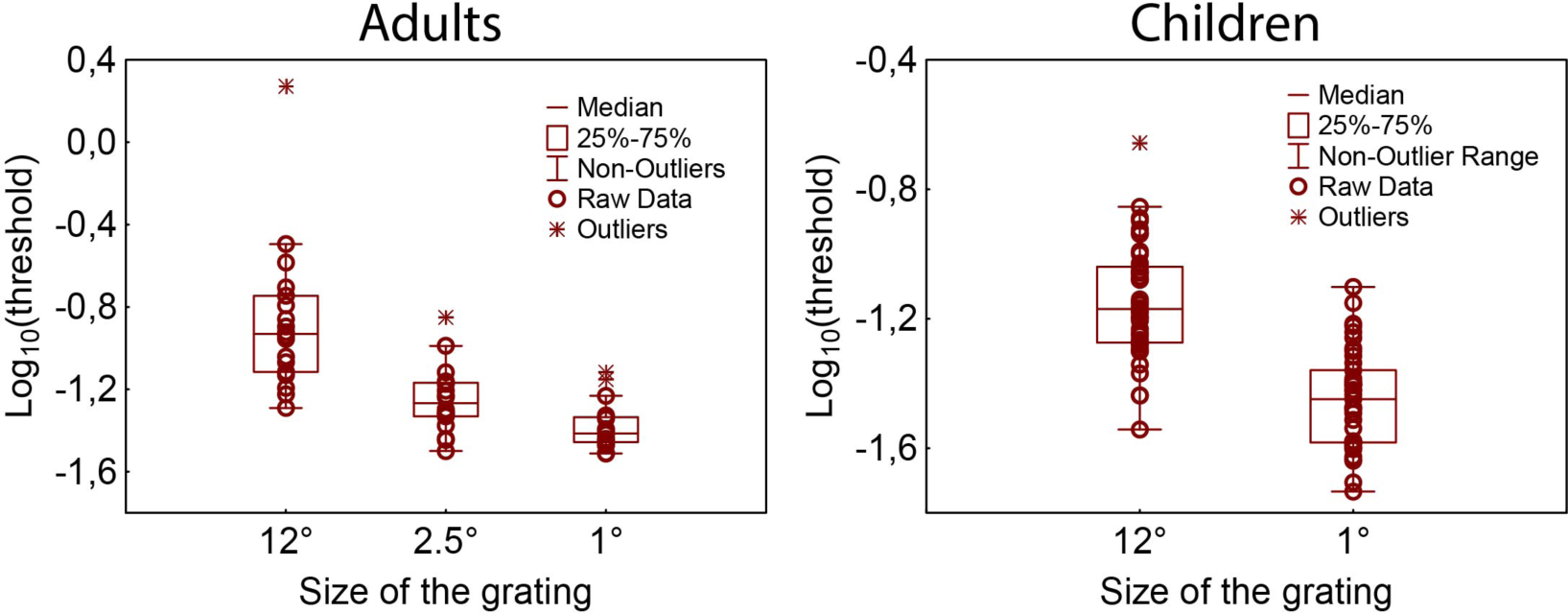
Log_10_ of direction discrimination thresholds (in seconds), measured in the psychophysical experiment in adults and children.

#### 3.1.2. Test-retest reliability of behavioral parameters in adults

In adults, we assessed test-retest reliability by measuring psychophysical thresholds in two sessions, separated by 5-118 days. The intra-class correlations between the two measurements are shown in table 1. The reliability was remarkably high for motion direction discrimination thresholds for the large gratings, somewhat lower for the small gratings, (insignificant for the medium gratings), and only moderate for the SSI. For the following analysis, we averaged the psychophysical measures across the two sessions.

**Table 1.** Test-retest reliability of psychophysical parameters in adults: intra-class correlations (ICC).

| Parameter | ICC |
| --- | --- |
| *Motion direction discrimination thresholds:* |  |
| Large | 0.95*** |
| Medium | 0.43 |
| Small | 0.75** |
| SSI | 0.51# |

#p<0.1, \*\*p<0.01; \*\*\*p<0.001

#### 3.1.3. Spatial suppression effect in children

Similar to adults, children needed significantly more time to discriminate the direction of motion of the large, as compared to the small, gratings (Chi-Sqr_(43, 1)_=35.4, p<0.0001, Kendel’s W=0.82; Fig. 3), that is indicative of reliable SS. The median thresholds were 36 ms for the small and 68 ms for the large gratings that corresponded to an approximately 100% rise in thresholds as the stimulus size increased from 1 to 12° of visual angle. The individual SS values ranged from −0.093 to 0.86 (−19% to 619% change in the duration threshold). Slight decreases in thresholds with increased stimulus size were observed in only 2 of 34 participants and could be explained by difficulties with reliable assessment of the psychophysical thresholds for these children.

### 3.2. MEG

#### 3.2.1. MEG gamma responses in adults

All of the adult participants demonstrated highly statistically reliable GRs (i.e. p<0.0001, see Materials and Methods for details on estimation of inter-individual reliability) during both measurement sessions. In source space, reliable GRs were observed in all adult participants for all four types of stimuli. In the majority of individual cases, as well as at the group level (Fig. 4A), the maximal gamma increase was localized to the calcarine fissure (area V1). In this respect, our data replicates the results of previous MEG studies that implicate the primary visual cortex as a generator of gamma oscillations induced by visual gratings (Adjamian et al., 2004; Hadjipapas et al., 2007; Hall et al., 2005; Hoogenboom et al., 2006; Muthukumaraswamy et al., 2010; Orekhova et al., 2019). Figures 4B and 4C show grand average time-frequency plots and spectra of the stimulus-related changes at the ‘maximal pair’ of gradiometers in adults and children.

**Fig. 4.**
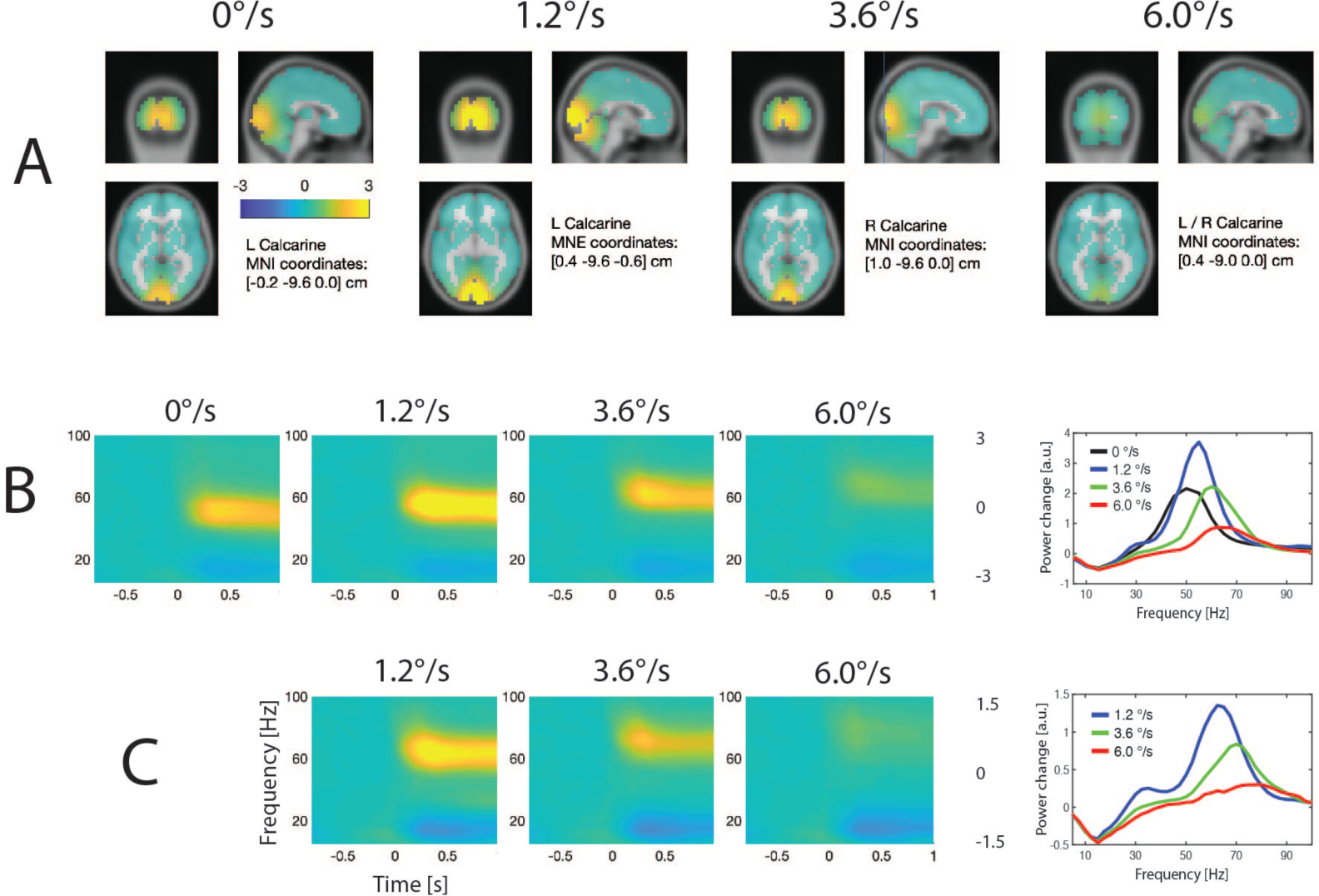
Spectral MEG changes induced by annular gratings drifting at different velocities. **A.** Source localization of the group average GR power in adults. The MNI coordinates correspond to a voxel where the maximal GR was observed at a group level. **B, C.** Grand average time-frequency plots and the corresponding spectra of the stimulation-related changes in MEG power at the ‘maximal pair’ of gradiometers in adults (B) and children (C). Colors correspond to velocity conditions: black – ‘static’, blue – ‘slow’, green – ‘medium’, red – ‘fast’.

In accordance with our previous results (Orekhova et al., 2015; Orekhova et al., 2019; Orekhova et al., 2018b), the GR frequency increased with increasing motion velocity from 0 to 6°/s (in source space: F_(3, 51)_ =97.8, p<1e-13; average frequency for static grating: 50.7 Hz, and for those drifting at different velocities - slow: 55.4 Hz, medium: 61.8 Hz, fast: 65.4 Hz). In the majority of the participants (15 of 18), the GR power increased with the transition from static to slowly moving stimuli and then decreased with increasing motion velocity (Fig. 5A), which is also consistent with our previous results (Orekhova et al., 2018b). This effect was highly reliable both in sensor (Chi-Sqr_(18, 3)_=40.2, p < 0.0001) and source (Chi-Sqr_(18, 3)_= 37.1, p < 0.0001) spaces.

**Fig. 5.**
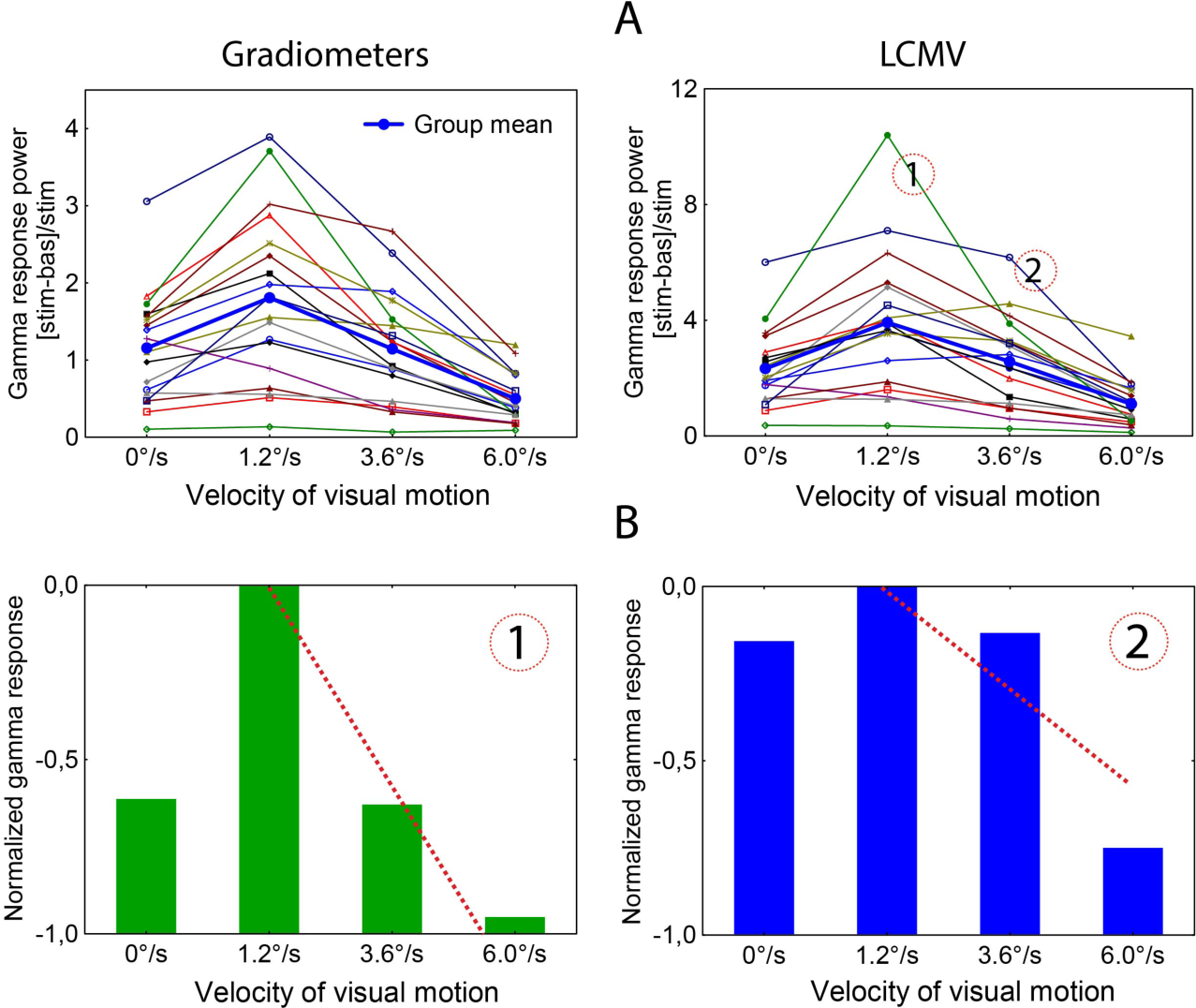
Dependence of GR power on the velocity of visual motion. **A.** Individual values of the GR power in adults, assessed in the sensor and source spaces. Note that after the initial increase from static to slowly moving gratings, the GR power decreases with increasing velocity of visual motion. **B.** The gamma suppression slope (GSS) – a measure that quantifies the decline in GR power as a function of visual motion velocity. Subject ‘1’ (marked ‘1’ in A) has strong GR attenuation, while subject ‘2’ (marked ‘2’ in A) has weaker.

In line with the previous reports (Muthukumaraswamy et al., 2010; Tan et al., 2016; van Pelt et al., 2012), there was considerable inter-individual variability of the GR power and frequency. As an example, when measured in source space, the power of the GR elicited by slowly moving gratings varied from subject to subject from 35% to 1040 % (relative to baseline) (Fig. 5A) and gamma frequency varied from 39 to 60 Hz.

We then analysed GSS – a measure of GR attenuation with increasing motion velocity for all individuals (Fig. 5B, see Materials and Methods for details). Despite considerable inter-individual variability, GSS was negative in all our participants, i.e. the power of the GR attenuated with increasing motion velocity from 1.2 to 6.0°/s in all subjects (Fig. 5A).

#### 3.2.2. Test/re-test reliability of gamma responses in adults

While the frequency and amplitude of visual gamma oscillations induced by static or slowly moving (1.6°/s) gratings are known to be highly individually stable traits (Hoogenboom et al., 2006; Muthukumaraswamy et al., 2010), the reliability of GSS – the measure of velocity-related attenuation of GR - has not been previously evaluated. In adults, we calculated the intra-class correlations for the gamma parameters measured at the two sessions separated by 5-118 days (mean=37.4 days, sd= 32.2, range 6-119; Table 2). Consistent with previous findings, MEG gamma frequency and power were found to be highly reliable measures over the temporal interval studied. The reliability of GSS was also high, especially when estimated in source space.

**Table 2.** Test/re-rest reliability of MEG gamma parameters in adults: intra-class correlations (ICC).

| Parameter | ICC |
| --- | --- |
| *MEG / GR frequency:* |  |
| 'Static' | 0.88*** (0.74**) |
| 'Slow velocity' | 0.93*** (0.97***) |
| 'Medium velocity' | 0.94*** (0.85**) |
| 'Fast velocity' | 0.81*** (0.74**) |
| *MEG / GR power:* |  |
| 'Static' | 0.94*** (0.87***) |
| 'Slow velocity' | 0.95*** (0.85***) |
| 'Medium velocity' | 0.88*** (0.74**) |
| 'Fast velocity' | 0.9*** (0.72**) |
| *gamma suppression slope (GSS)* | 0.91*** (0.77**) |

Note: the correlations are given for source and sensor (in parentheses) spaces; \*\*p<0.01; \*\*\*p<0.001

#### 3.2.3. MEG gamma responses in children

We have described the effects of visual motion velocity on gamma parameters in children elsewhere (Orekhova et al., 2018b). Here, these effects were reproduced in the smaller sample of participants included in the current study and were similar to those found in adults. Specifically, the GR power decreased (ANOVA Chi Sqr_(34, 2)_ = 60, p < 0.0001), while the GR frequency increased (Chi Sqr _(19, 2)_ = 38, p < 0.0001), with increasing velocity of visual motion from 1.2 to 6 °/s (Fig. 4C). Similarly to adults, all children with reliable GRs in the ‘slow’ condition had negative GSS.

### 3.3. Gamma suppression slope and spatial suppression

#### 3.3.1. Adults

As predicted, there was a significant correlation between SS and velocity-related GR attenuation: participants with stronger GR attenuation (i.e. more negative GSS) had higher SSI (SSI vs GSS in sensor space: R_(18)_=-0.51, p<0.05, in source space: R_(18)_=-0.71, p<0.001, Fig. 6). We then looked at the correlations between GSS and the motion direction discrimination thresholds for large and small gratings. A stronger GR attenuation was associated with more time needed for a subject to discriminate the motion direction of the large grating (in sensor space: R_(18)_=-0.63, p<0.01; in source space: R_(18)_=-0.64, p<0.01, see also Fig. 6, lower left panel). Exclusion of the outlier with an extremely high motion direction discrimination threshold for the large grating (Fig. 3, left panel) did not significantly affect the results (SSI vs GSS in sensor space: R_(17)_=-0.48, p=0.05; in source space: R_(17)_= −0.69, p<0.01; Threshold_LARGE_ vs GSS in sensor space: R_(17)_=-0.61, p<0.01; in source space: R_(17)_=-0.61, p<0.01). Concurrently, the thresholds for the small or medium gratings did not correlate with GSS (GSS in sensor space vs Threshold_SMALL_: R_(18)_=-0.13 vs Threshold_MEDIUM_: R_(18)_=- 0.34; GSS in source space vs Threshold_SMALL_: R_(18)_=0.014, Threshold_MEDIUM_: R_(18)_=-0.06). Thus, GSS magnitude appears to be selectively related to a subject’s capacity to detect the motion direction of the large gratings.

**Fig. 6.**
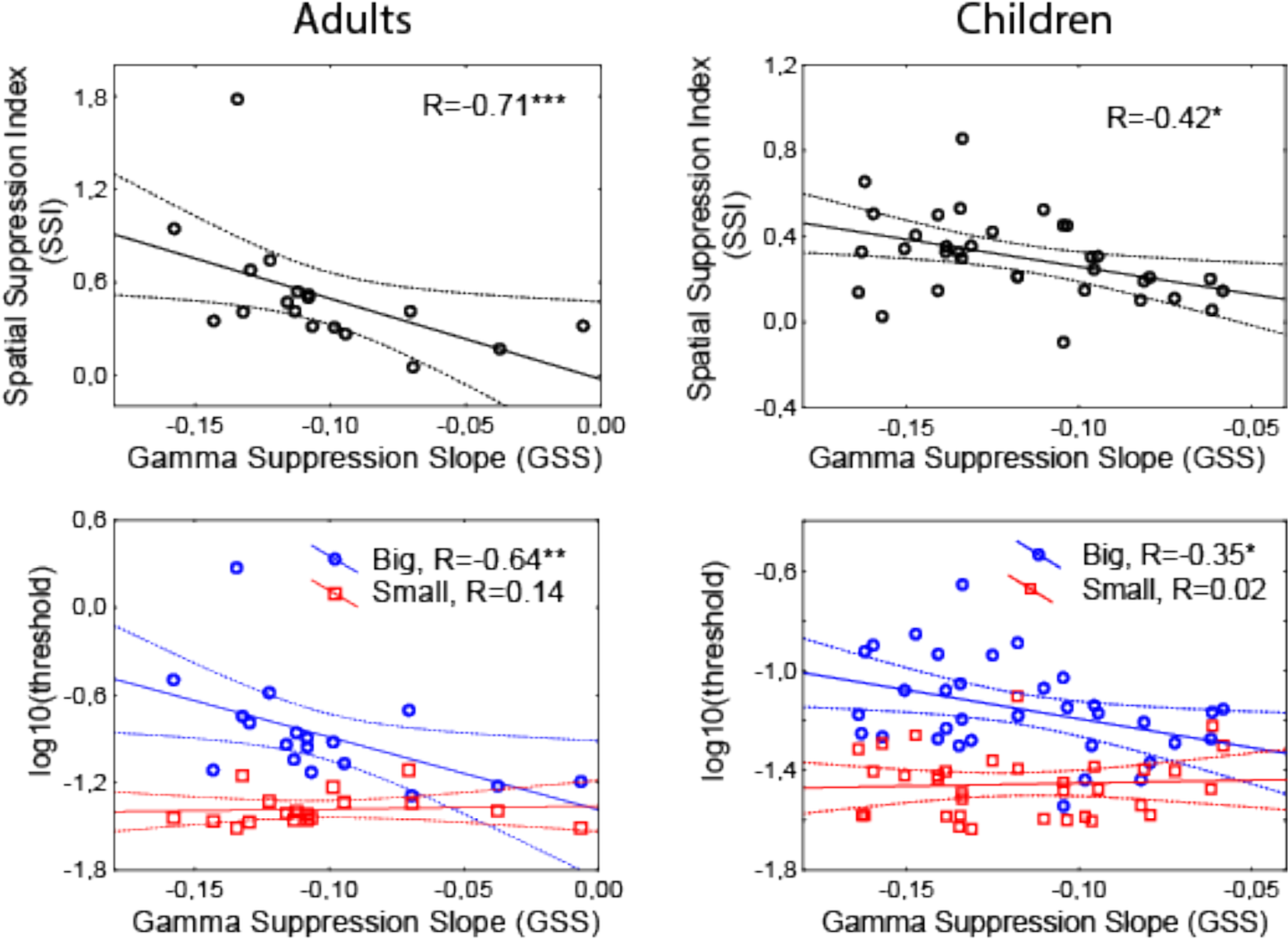
The relationship between GR suppression and SS in adults and children. **Upper row**: relationship between GSS and SSI. **Lower row:** the relationship between GSS and duration thresholds. Note that no reliable correlations were found between GSS and duration thresholds for the small stimuli. GSS is shown for the brain sources in adults and for the pairs of MEG sensors in children. Solid lines denote the regression; dashed lines mark 95% confidence intervals. R’s are the Spearman correlation coefficients. *p<0.05, **p<0.01, ***p<0.001.

To examine whether SSI is specifically linked to GSS, we correlated this psychophysical index with other parameters of the GR. Neither gamma peak frequency nor gamma peak power predicted SSI; neither of them predicted the motion direction discrimination threshold for the large grating, either (Spearman R’s: all p’s>0.1 for all velocities in both sensor and source spaces). In particular, absence of significant correlations between SSI and GR magnitudes indicates that the correlation between SSI and GSS was not driven by GR magnitude under any specific velocity condition.

#### 3.3.2. Test/re-test reliability of neurobehavioral correlations in adults

To ensure that the relationship between GSS and psychophysical parameters was reproducible across time, we calculated the neuro-behavioral correlations for the measurements obtained at the 1^st^ and the 2^nd^ visits. There was good agreement in these correlations between the visits (In source space: 1^st^ visit GSS vs SSI: R_(18)_=-0.58, p<0.05, GSS vs Threshold_BIG_: R_(18)_=-0.57, p<0.05; 2^nd^ visit GSS vs SSI: R_(18)_=-0.53, p<0.05, GSS vs Threshold_BIG_: R_(18)_=-0.60, p<0.01).

#### 3.3.3. Children

Similarly to adults, stronger SS was associated with more negative GSS measured in the sensor space (R_(34)_=-0.42, p<0.05; Fig. 6). In other words, those children who had stronger SS also demonstrated stronger velocity-related GR attenuation. Also, in striking similarity to adults, correlation with GSS in children was driven by the thresholds for the large gratings (R_(34)_=-0.35, p<0.05; Fig. 6), while no reliable correlation was found for the small grating (R_(34)_=0.02).

Overall, the observed relationship between velocity-related GR attenuation and SS matches the hypothesis of their common inhibitory basis and suggest its generality across the genders and ages studied.

### 3.4. Gamma suppression slope, spatial suppression and IQ

Previous studies have suggested that stronger inhibitory control in the visual cortex favors efficient cognitive functioning (Cook et al., 2016; Melnick et al., 2013). Therefore, the strength of inhibition measured through GSS may affect not only SS, but also contribute to higher cognitive functions. Indeed, we found that children with more negative GSS had higher IQ (KABC-II, Mental Processing Index (MPI): R_(34)_=0.41, p<0.05 (Fig. 7). At the same time, SS strength did not correlate with IQ (R _(34)_ =-0.04). Information about IQ was not available for the adult participants; generalization of these findings to adults is thus not possible.

**Fig. 7.**
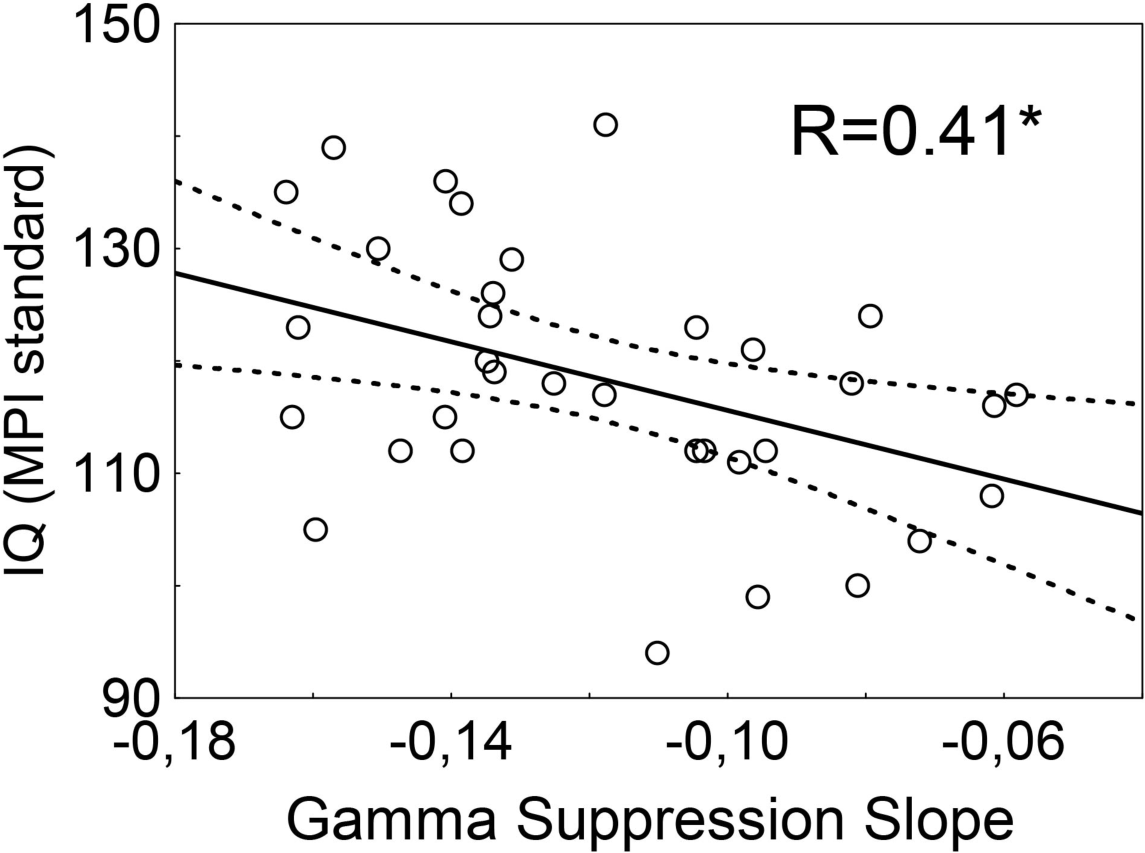
The relationship between GSS and IQ in children. Solid lines denote the regression; dashed lines mark 95% confidence intervals. R is the Spearman correlation coefficient; *p<0.05

## 4. Discussion

Using a behavioral task and magnetoencephalography, we tested for a link between two phenomena, both of which have been suggested to depend on efficacy of inhibitory control in the visual cortex. The first one – SS in visual motion perception - refers to the reduction of directional sensitivity to large, as compared to small, rapidly moving high-contrast visual gratings. The second one – GSS – is a progressive attenuation of MEG GR with increasing velocity of the large high-contrast visual gratings. We found that a stronger GR attenuation predicted stronger SS, i.e. greater deterioration of behavioral performance with increasing size of the grating. This neuro-behavioral correlation was replicated in two independent samples of participants – adult women and boys aged 7-15 years. Neither GR peak frequency nor its peak power measured in a single experimental condition predicted SS in any group of subjects. Our findings therefore highlight the functional relevance of velocity-related *changes* in GR power caused by augmentation of incoming input. In the following discussion, we consider a possible neural basis for the observed neuro-behavioral correlation.

### 4.1. SS, MEG gamma oscillations, and surround suppression in the visual cortex

The presence of a reliable SS is a necessary prerequisite for finding its correlation with MEG gamma-based measures. The SS effect assessed in a separate psychophysical experiment in our study was reliable in both adults and children: the participants needed significantly longer presentation times in order to discriminate motion direction of the large, as compared to small, visual gratings (Fig. 2). Moreover, our assessment of the test-retest reliability of direction discrimination thresholds performed in adults has shown that it was high (Table 1).

In accordance with our previously published results (Orekhova et al., 2015; Orekhova et al., 2018a), large high-contrast visual gratings, when presented at the ‘optimal’ velocity of 1.2°/s, induced strong GRs in all adult participants and in the majority of children. Also in accordance with previous studies (Adjamian et al., 2004; Hadjipapas et al., 2007; Hall et al., 2005; Hoogenboom et al., 2006; Muthukumaraswamy et al., 2010; Orekhova et al., 2015; Orekhova et al., 2018a), the maximum of the visual GR was located in V1. The combination of regular orientation, high contrast, large size, and slow motion velocity of the grating facilitated high amplitude gamma oscillations. Indeed, each of these visual features enhances GRs in the primary visual cortex in humans and animals (Gieselmann and Thiele, 2008; Hadjipapas et al., 2015; Hermes et al., 2015; Jia et al., 2011; Jia et al., 2013; Lima et al., 2010; Orekhova et al., 2015; Orekhova et al., 2018a; Perry, 2015; Perry et al., 2013; Perry et al., 2015; Ray and Maunsell, 2015; van Pelt and Fries, 2013).

The crucial factor affecting visual gamma oscillations seems to be the size of the visual stimulus --- gamma is generated only when the stimulus is large enough to engage both the center and the inhibitory surround of the neurons’ receptive fields (Gieselmann and Thiele, 2008; Shushruth et al., 2012). Gieselmann and Thiele (Gieselmann and Thiele, 2008) have shown that local field potential (LFP) gamma-band activity increased monotonically with grating size, despite a concomitant drop in neuronal spiking rates. The increase in LFP gamma power was maximal for stimuli overlapping the near classical receptive field surround, where suppression started to dominate spiking activity (Gieselmann and Thiele, 2008).

The large annular gratings used in our MEG experiment (18° of visual angle) should have led to the strong involvement of the inhibitory surround in the populational neural activity at the V1 level. Therefore, the changes in GR elicited by increasing the gratings’ drift rate can be a direct consequence of surround modulation effects in V1. A nearly linear growth of GR frequency with increasing visual motion velocity from 0 up to 6°/s in our study most likely reflects a growing of intensity in the excitatory drive and related growth of excitability of excitatory, and especially inhibitory neurons; the later being largely responsible for changes in frequency of gamma oscillations (Anver et al., 2011; Mann and Mody, 2010). The same factor may account for the inverted parabolic trajectory of the GR power (Fig. 5; see (Orekhova et al., 2019) for discussion). In particular, the reduction of the GR with increasing motion velocity beyond the ‘optimal’ value of ~1.2°/s is likely to be driven by an excessively strong involvement of the inhibitory surround in the populational neural activity at the V1 level. In our present study, the suppression of the visual GR at motion velocity above 1.2-3.6°/s has been observed in every subject, irrespectively of his/her age or gender, suggesting that the velocity-related gamma attenuation reflects this basic inhibitory-based mechanism of cortical function.

MEG is capable of detecting gamma oscillations only if the coherent activity in the gamma range is observed in a large portion of the visual cortex. Therefore, the presence of prominent GRs in our MEG study suggests widespread in-phase gamma synchronization in the visual cortex. Indeed, animal studies have revealed phase-alignment of LFP gamma oscillations over a broad V1 area (Eckhorn et al., 1988; Jia et al., 2011). One plausible mechanism of such distributed gamma synchronization is feedback connections from the higher-tier visual areas, which extend over long distances and play a pivotal role in surround suppression (Angelucci et al., 2017; Angelucci and Bressloff, 2006; Nurminen et al., 2018) as well as its perceptual consequence i.e., SS (Liu et al., 2016; Tadin, 2015; Tadin et al., 2003). Being strongly engaged by spatially extended fast moving stimuli, the higher-tier visual areas, including MT, may affect both SS and GR through potentiating surround suppression in V1.

The role of surround inhibition in both SS and GR power is supported by the key finding of the current study replicated in the two independent samples of subjects of different age and gender (Fig. 6): the psychophysically measured SSI significantly correlates with the GSS index (the latter of which quantifies the attenuation of GR in V1 at high velocities of visual motion). As predicted, in both samples of subjects, GSS also correlated with duration thresholds for the *large* moving grating that causes surround inhibition, but not with the same for the small grating that does not. This selectivity strongly implicates surround inhibition in the observed neuro-behavioral correlation.

The hypothesis that intensive visual input (e.g. gratings that are large in size, high in contrast, and moving with high velocity) is a factor that mediates GSS-SSI correlation through a disproportionate increase in surround inhibition is compatible with theoretical models of surround inhibition (Rubin et al., 2015). Recent neurophysiological evidence furthermore directly verifies these models (Adesnik, 2017). Adesnik and colleagues performed intracellular voltage-clamp measurements of excitatory and inhibitory post-synaptic currents in the primary visual cortex of awake mice and found that increasing the size and/or contrast of a visual grating leads to a gradual increase in excitatory current paralleled by more steep increase of inhibitory current; the result is thus a decline in the E/I ratio.

Our findings, as well as the results of animal studies, seem to conflict with the ‘withdrawal of excitation’ explanation for SS put forth by Schallmo et al (Schallmo et al., 2018). Those authors concluded that surround suppression in both V1 and the higher-tier visual area MT is primarily driven by a decrease in excitation, which is balanced by reduced inhibition in visual networks (Schallmo et al., 2018). A lack of significant correlations between GABA concentration at rest and either psychophysically measured SS or the size-dependent decrease in BOLD signal in MT and V1 regions supports this explanation. However, inter-individual differences in neural inhibition may depend on complex interactions between numerous pre-synaptic, post-synaptic, and extra-synaptic mechanisms (Isaacson and Scanziani, 2011) and might not be reflected in MRS measurements of GABA (Goncalves et al., 2017). The ‘withdrawal of excitation’ hypothesis furthermore fails to provide meaningful explanation for the velocity dependence of the GR frequency in the face of experimental results. Large gratings moving at velocities greater than 1°/s produce stronger spatial suppression than those moving at slower velocities (Lappin et al., 2009). Faster velocities within the 0°/s – 6 °/s range furthermore produce a substantial increase in visual GR frequency (Fig. 4; see also van Pelt et al, 2018). Such an increase indicates excitation is *elevated* in inhibitory neurons in human V1 because the GR frequency directly depends on the excitatory state of inhibitory neurons (Anver et al., 2011; Mann and Mody, 2010). However, if SS is driven by ‘withdrawal of excitation’, then faster motion velocities should presumably lead to *reduced* excitation of both excitatory and inhibitory neurons in V1. Because such a reduction in excitation should be accompanied by a decrease in gamma response frequency, this presumption is contrary to the evidence. On the other hand, the strong size-dependent reduction in the V1 BOLD signal, paralleled by a less pronounced reduction in MT+, reported by Schallmo et al (Schallmo et al., 2018) does not contradict the model suggested by Angelucci (Angelucci et al., 2017; Angelucci and Bressloff, 2006), which posits that SS is a direct consequence of center– surround antagonism within V1, while MT enhances this antagonism by potentiating inhibition in V1 through providing excitatory feedback to V1.

### 4.2. Replication of the results and test-retest reliability

An important advantage of our study is the testing for replication of the anticipated neuro-behavioral correlations in two diverse groups of participants, as well as estimation of test-retest reliability of these correlations and the neurophysiological and psychophysical parameters themselves.

First, the correlation between GSS and SS was replicated in the two groups of subjects i.e., GSS correlates with SS in both adult women and male children (Fig. 6). This suggests the results are robust with respect to sampling bias and can be generalized across gender and ages studied.

Second, the MEG gamma characteristics and psychophysical thresholds were replicated in the two temporally-separated measurement sessions i.e., we report a high test-retest reliability of these parameters in sessions that were separated by several days or weeks. The high test-retest reliability of GSS is, at first glance, similar to that of the GR power and frequency evaluated separately in each velocity condition (Table 2; see also (Hoogenboom et al., 2006; Muthukumaraswamy et al., 2010). However, constitutional factors including cortical morphology (Butler et al., 2019) and degree of muscle tension (Muthukumaraswamy, 2013; Whithain et al., 2007; Whitham et al., 2008) can specifically affect MEG/EEG gamma power while gamma frequency may be affected by the size of the V1 area (Schwarzkopf et al., 2012; van Pelt et al., 2018). These features, that are not directly related to the functioning of neural networks, contribute to the estimated intra-individual stability of gamma frequency and amplitude measured under a single experimental condition. However, and in contrast to gamma power and frequency, GSS is a relational measure that relies upon between-stimulus differences in response strength, which are not (or less) affected by brain morphology and SNR. Therefore, individual differences in GSS are likely to be trait-like and mainly of neurophysiological origin.

### 4.3. SS, MEG gamma oscillations, and IQ in children

A collateral, but remarkable finding is the significant correlation between GSS and psychometric intelligence in children, for whom IQ information was available. Notably, there were no correlations between IQ and either SSI or motion direction discrimination thresholds. If individual variations in both GSS and SSI reflect variability in neural mechanisms that are responsible for surround suppression, why would IQ correlate with only one of these measures, namely GSS? Of note, there are previous reports of strong (Melnick et al., 2013) or moderate (Arranz-Paraiso and Serrano-Pedraza, 2018) SSI-IQ correlations in adults. However, the recent large (177 participants) and methodologically advanced study by Troche et al failed to replicate this SSI-IQ association (Troche et al., 2018). One of the reasons for these divergent results might be the fact that participants with higher intelligence were faster to detect motion direction for all (large and small) stimuli (Troche et al., 2018). This dependency of IQ on processing speed may obscure the IQ-SSI correlation, which, when found, showed that duration thresholds of motion direction discrimination for large and small stimuli correlate with IQ in opposite directions (Melnick et al., 2013). Specifically, higher IQ was associated with higher duration thresholds for large, but lower duration thresholds for the small stimuli (Melnick et al., 2013).

Thus, beyond individual differences in surround suppression, SSI depends on other psychological variables that might affect the results of the psychophysical experiment. Neurophysiological variables, such as GSS, are less likely to depend on processing speed, and may thus reflect inter-individual differences in surround suppression better than psychophysical measures do. If GSS indeed reflects inhibition efficiency, its correlation with IQ is well in line with the role of ‘inhibition-stabilized networks’ in cortical computations (Rubin et al., 2015). Moreover, the efficiency of recurrent feedback to V1 from higher-tier visual areas (Angelucci et al., 2017; Angelucci and Bressloff, 2006; Nurminen et al., 2018) may reflect general efficacy of top-down modulations important for cognitive control and contributing to the link between IQ and neural gamma dynamics in the primary visual cortex.

*To summarize*, our study provides the first evidence that links the psychophysical phenomenon of spatial suppression and the inhibitory-based neural gamma responses in the human primary visual cortex. The results are replicated in two groups of subjects - adult women and male children, which suggests they are robust and general across age and gender. Our findings support the role of inhibitory interactions in the V1 as the mechanism of spatial suppression.

## Acknowledgements

This work has been supported by the research grant from The Moscow State University of Psychology and Education (MSUPE), The Knut and Alice Wallenberg foundation (KAW2014.0102) and The Swedish Childhood Cancer Foundation (MT2018-0020).

## References

Adesnik, H., 2017. Synaptic Mechanisms of Feature Coding in the Visual Cortex of Awake Mice. Neuron 95, 1147–1159.

Adjamian, P., Holliday, I.E., Barnes, G.R., Hillebrand, A., Hadjipapas, A., Singh, K.D., 2004. Induced visual illusions and gamma oscillations in human primary visual cortex. Eur J Neurosci 20, 587–592.

Angelucci, A., Bijanzadeh, M., Nurminen, L., Federer, F., Merlin, S., Bressloff, P.C., 2017. Circuits and Mechanisms for Surround Modulation in Visual Cortex. Annual Review of Neuroscience, Vol 40 40, 425–451.

Angelucci, A., Bressloff, P.C., 2006. Contribution of feedforward, lateral and feedback connections to the classical receptive field center and extra-classical receptive field surround of primate V1 neurons. Prog Brain Res 154, 93–120.

Anver, H., Ward, P.D., Magony, A., Vreugdenhil, M., 2011. NMDA Receptor Hypofunction Phase Couples Independent gamma-Oscillations in the Rat Visual Cortex. Neuropsychopharmacology 36, 519–528.

Arranz-Paraiso, S., Serrano-Pedraza, I., 2018. Testing the link between visual suppression and intelligence. Plos One.

Atallah, B.V., Bruns, W., Carandini, M., Scanziani, M., 2012. Parvalbumin-Expressing Interneurons Linearly Transform Cortical Responses to Visual Stimuli. Neuron 73, 159–170.

Baddeley, R., Abbott, L.F., Booth, M.C.A., Sengpiel, F., Freeman, T., Wakeman, E.A., Rolls, E.T., 1997. Responses of neurons in primary and inferior temporal visual cortices to natural scenes. Proceedings of the Royal Society B-Biological Sciences 264, 1775–1783.

Barlow, H.B., 1961. The coding of sensory messages. In: Thorpe, W.H., Zangwill, L. (Eds.), Current Problems in Animal Behavior. Cambridge U. Press, Cambridge, pp. 331–360.

Betts, L.R., Sekuler, A.B., Bennett, P.J., 2007. The effects of aging on orientation discrimination. Vision Research 47, 1769–1780.

Betts, L.R., Sekuler, A.B., Bennett, P.J., 2012. Spatial characteristics of motion-sensitive mechanisms change with age and stimulus spatial frequency. Vision Research 53, 1–14.

Borgers, C., Kopell, N., 2005. Effects of noisy drive on rhythms in networks of excitatory and inhibitory neurons. Neural Computation 17, 557–608.

Born, R.T., Bradley, D.C., 2005. Structure and function of visual area MT. Annual Review of Neuroscience, Vol 40 28, 157–189.

Brainard, D.H., 1997. The psychophysics toolbox. Spat. Vis. 10, 433–436.

Butler, R., Bernier, P.M., Mierzwinski, G.W., Descoteaux, M., Gilbert, G., Whittingstall, K., 2019. Cortical distance, not cancellation, dominates inter-subject EEG gamma rhythm amplitude. Neuroimage 192, 156–165.

Button, K.S., Ioannidis, J.P.A., Mokrysz, C., Nosek, B.A., Flint, J., Robinson, E.S.J., Munafo, M.R., 2013. Power failure: why small sample size undermines the reliability of neuroscience. Nat Rev Neurosci 14, 365–376.

Buzsaki, G., Anastassiou, C.A., Koch, C., 2012. The origin of extracellular fields and currents - EEG, ECoG, LFP and spikes. Nat Rev Neurosci 13, 407–420.

Buzsaki, G., Wang, X.J., 2012. Mechanisms of Gamma Oscillations. Annual Review of Neuroscience, Vol 40 35, 203–225.

Cannon, J., McCarthy, M.M., Lee, S., Lee, J., Borgers, C., Whittington, M.A., Kopell, N., 2014. Neurosystems: brain rhythms and cognitive processing. Eur J Neurosci 39, 705–719.

Cardin, J.A., Carlen, M., Meletis, K., Knoblich, U., Zhang, F., Deisseroth, K., Tsai, L.H., Moore, C.I., 2009. Driving fast-spiking cells induces gamma rhythm and controls sensory responses. Nature 459, 663–U663.

Cook, E., Hammett, S.T., Larsson, J., 2016. GABA predicts visual intelligence. Neurosci Lett 632, 50–54.

Desimone, R., Duncan, J., 1995. Neural mechanisms of selective visual attention. Annual Review of Neuroscience, Vol 40 18, 193–222.

Eckhorn, R., Bauer, R., Jordan, W., Brosch, M., Kruse, W., Munk, M., Reitboeck, H.J., 1988. Coherent Oscillations - a Mechanism of Feature Linking in the Visual-Cortex - Multiple Electrode and Correlation Analyses in the Cat. Biological Cybernetics 60, 121–130.

Field, D.J., 1987. Relations between the Statistics of Natural Images and the Response Properties of Cortical-Cells. Journal of the Optical Society of America a-Optics Image Science and Vision 4, 2379–2394.

Foss-Feig, J.H., Tadin, D., Schauder, K.B., Cascio, C.J., 2013. A Substantial and Unexpected Enhancement of Motion Perception in Autism. J Neurosci 33, 8243–8249.

Gieselmann, M.A., Thiele, A., 2008. Comparison of spatial integration and surround suppression characteristics in spiking activity and the local field potential in macaque V1. Eur J Neurosci 28, 447–459.

Golomb, J.D., McDavitt, J.R.B., Ruf, B.M., Chen, J.I., Saricicek, A., Maloney, K.H., Hu, J., Chun, M.M., Bhagwagar, Z., 2009. Enhanced Visual Motion Perception in Major Depressive Disorder. J Neurosci 29, 9072–9077.

Goncalves, J., Violante, I.R., Sereno, J., Leitao, R.A., Cai, Y., Abrunhosa, A., Silva, A.P., Silva, A.J., Castelo-Branco, M., 2017. Testing the excitation/inhibition imbalance hypothesis in a mouse model of the autism spectrum disorder: in vivo neurospectroscopy and molecular evidence for regional phenotypes. Mol Autism 8, 47.

Gramfort, A., Luessi, M., Larson, E., Engemann, D.A., Strohmeier, D., Brodbeck, C., Goj, R., Jas, M., Brooks, T., Parkkonen, L., Hamalainen, M., 2013. MEG and EEG data analysis with MNE-Python. Front Neurosci.

Hadjipapas, A., Adjamian, P., Swettenham, J.B., Holliday, I.E., Barnes, G.R., 2007. Stimuli of varying spatial scale induce gamma activity with distinct temporal characteristics in human visual cortex. Neuroimage 35, 518–530.

Hadjipapas, A., Lowet, E., Roberts, M.J., Peter, A., De Weerd, P., 2015. Parametric variation of gamma frequency and power with luminance contrast: A comparative study of human MEG and monkey LFP and spike responses. Neuroimage 112, 327–340.

Hall, S.D., Holliday, I.E., Hillebrand, A., Singh, K.D., Furlong, P.L., Hadjipapas, A., Barnes, G.R., 2005. The missing link: analogous human and primate cortical gamma oscillations. Neuroimage 26, 13–17.

Hasenstaub, A., Shu, Y.S., Haider, B., Kraushaar, U., Duque, A., McCormick, D.A., 2005. Inhibitory postsynaptic potentials carry synchronized frequency information in active cortical networks. Neuron 47, 423–435.

Hermes, D., Miller, K.J., Wandell, B.A., Winawer, J., 2015. Stimulus Dependence of Gamma Oscillations in Human Visual Cortex. Cerebral Cortex 25, 2951–2959.

Hoogenboom, N., Schoffelen, J.M., Oostenveld, R., Parkes, L.M., Fries, P., 2006. Localizing human visual gamma-band activity in frequency, time and space. Neuroimage 29, 764–773.

Isaacson, J.S., Scanziani, M., 2011. How Inhibition Shapes Cortical Activity. Neuron 72, 231–243.

Jia, X.X., Smith, M.A., Kohn, A., 2011. Stimulus Selectivity and Spatial Coherence of Gamma Components of the Local Field Potential. J Neurosci 31, 9390–9403.

Jia, X.X., Xing, D.J., Kohn, A., 2013. No Consistent Relationship between Gamma Power and Peak Frequency in Macaque Primary Visual Cortex. J Neurosci 33, 17–U421.

Kaufman, A.S., Kaufman, N.L., 2004. KABC-II: Kaufman Assessment Battery for Children, 2nd ed., 2 ed. AGS Pub, Circle Pines, MN.

King, P.D., Zylberberg, J., DeWeese, M.R., 2013. Inhibitory interneurons decorrelate excitatory cells to drive sparse code formation in a spiking model of V1. J Neurosci 33, 5475–5485.

Kopell, N., Börgers, C., Pervouchine, D., Malerba, P., Tort, A., 2010. Gamma and Theta Rhythms in Biophysical Models of Hippocampal Circuits. Springer Series in Computational Neuroscience Springer Science+Business Media.

Lappin, J.S., Tadin, D., Nyquist, J.B., Corn, A.L., 2009. Spatial and temporal limits of motion perception across variations in speed, eccentricity, and low vision. Journal of Vision.

Lima, B., Singer, W., Chen, N.H., Neuenschwander, S., 2010. Synchronization Dynamics in Response to Plaid Stimuli in Monkey V1. Cerebral Cortex 20, 1556–1573.

Liu, L.D., Haefner, R.M., Pack, C.C., 2016. A neural basis for the spatial suppression of visual motion perception. Elife.

Mann, E.O., Mody, I., 2010. Control of hippocampal gamma oscillation frequency by tonic inhibition and excitation of interneurons. Nature Neuroscience 13, 205–U290.

Melnick, M.D., Harrison, B.R., Park, S., Bennetto, L., Tadin, D., 2013. A strong interactive link between sensory discriminations and intelligence. Curr Biol 23, 1013–1017.

Muthukumaraswamy, S.D., 2013. High-frequency brain activity and muscle artifacts in MEG/EEG: a review and recommendations. Frontiers in Human Neuroscience.

Muthukumaraswamy, S.D., Singh, K.D., Swettenham, J.B., Jones, D.K., 2010. Visual gamma oscillations and evoked responses: Variability, repeatability and structural MRI correlates. Neuroimage 49, 3349–3357.

Nurminen, L., Angelucci, A., 2014. Multiple components of surround modulation in primary visual cortex: multiple neural circuits with multiple functions? Vision Research 104, 47–56.

Nurminen, L., Merlin, S., Bijanzadeh, M., Federer, F., Angelucci, A., 2018. Top-down feedback controls spatial summation and response amplitude in primate visual cortex. Nature Communications.

Olshausen, B.A., Field, D.J., 2004. Sparse coding of sensory inputs. Current Opinion in Neurobiology 14, 481–487.

Oostenveld, R., Fries, P., Maris, E., Schoffelen, J.M., 2011. FieldTrip: Open source software for advanced analysis of MEG, EEG, and invasive electrophysiological data. Comput Intell Neurosci.

Orekhova, E.V., Butorina, A.V., Sysoeva, O.V., Prokofyev, A.O., Nikolaeva, A.Y., Stroganova, T.A., 2015. Frequency of gamma oscillations in humans is modulated by velocity of visual motion. Journal of Neurophysiology 114, 244–255.

Orekhova, E.V., Prokofyev, A.O., Nikolaeva, A.Y., Schneiderman, J.F., Stroganova, T.A., 2018a. Additive effect of contrast and velocity proves the role of strong excitatory drive in suppression of visual gamma response. BioRxiv.

Orekhova, E.V., Stroganova, T.A., Schneiderman, J.F., Lundstrom, S., Riaz, B., Sarovic, D., Sysoeva, O.V., Brant, G., Gillberg, C., Hadjikhani, N., 2019. Neural gain control measured through cortical gamma oscillations is associated with sensory sensitivity. Hum Brain Mapp 40, 1583–1593.

Orekhova, E.V., Sysoeva, O.V., Schneiderman, J.F., Lundstrom, S., Galuta, I.A., Goiaeva, D.E., Prokofyev, A.O., Riaz, B., Keeler, C., Hadjikhani, N., Gillberg, C., Stroganova, T.A., 2018b. Input-dependent modulation of MEG gamma oscillations reflects gain control in the visual cortex. Sci Rep 8, 8451.

Perry, G., 2015. The effects of cross-orientation masking on the visual gamma response in humans. Eur J Neurosci 41, 1484–1495.

Perry, G., Hamandi, K., Brindley, L.M., Muthukumaraswamy, S.D., Singh, K.D., 2013. The properties of induced gamma oscillations in human visual cortex show individual variability in their dependence on stimulus size. Neuroimage 68, 83–92.

Perry, G., Randle, J.M., Koelewijn, L., Routley, B.C., Singh, K.D., 2015. Linear Tuning of Gamma Amplitude and Frequency to Luminance Contrast: Evidence from a Continuous Mapping Paradigm. Plos One.

Ray, S., Maunsell, J.H.R., 2015. Do gamma oscillations play a role in cerebral cortex? Trends in Cognitive Sciences 19, 78–85.

Rubin, D.B., Van Hooser, S.D., Miller, K.D., 2015. The Stabilized Supralinear Network: A Unifying Circuit Motif Underlying Multi-Input Integration in Sensory Cortex. Neuron 85, 402–417.

Sachdev, R.N.S., Krause, M.R., Mazer, J.A., 2012. Surround suppression and sparse coding in visual and barrel cortices. Frontiers in Neural Circuits.

Schallmo, M.P., Kale, A.M., Millin, R., Flevaris, A.V., Brkanac, Z., Edden, R.A., Bernier, R.A., Murray, S.O., 2018. Suppression and facilitation of human neural responses. Elife.

Schwarzkopf, D.S., Robertson, D.J., Song, C., Barnes, G.R., Rees, G., 2012. The frequency of visually induced gamma-band oscillations depends on the size of early human visual cortex. J Neurosci 32, 1507–1512.

Shushruth, S., Mangapathy, P., Ichida, J.M., Bressloff, P.C., Schwabe, L., Angelucci, A., 2012. Strong Recurrent Networks Compute the Orientation Tuning of Surround Modulation in the Primate Primary Visual Cortex. J Neurosci 32, 308–321.

Sohal, V.S., Zhang, F., Yizhar, O., Deisseroth, K., 2009. Parvalbumin neurons and gamma rhythms enhance cortical circuit performance. Nature 459, 698–702.

Stroganova, T.A., Sysoeva, O.V., Davletshina, M.S., Galuta, I.A., Goiaeva, D.E., Prokofyev, A.O., Orekhova, E.V., 2018. MEG Gamma Oscillations and Directional Sensitivity to Visual Motion in Children with ASD: Two Sides of the Inhibition Deficit. INSAR 2018 Annual Meeting. International Society for Autism Research, Rotterdam, Netherlands.

Sysoeva, O.V., Galuta, I.A., Davletshina, M.S., Orekhova, E.V., Stroganova, T.A., 2017. Abnormal Size-Dependent Modulation of Motion Perception in Children with Autism Spectrum Disorder (ASD). Front Neurosci 11, 164.

Szucs, D., Ioannidis, J.P.A., 2017. Empirical assessment of published effect sizes and power in the recent cognitive neuroscience and psychology literature. Plos Biology.

Tadin, D., 2015. Suppressive mechanisms in visual motion processing: From perception to intelligence. Vision Research 115, 58–70.

Tadin, D., Kim, J., Doop, M.L., Gibson, C., Lappin, J.S., Blake, R., Park, S., 2006. Weakened center-surround interactions in visual motion processing in schizophrenia. J Neurosci 26, 11403–11412.

Tadin, D., Lappin, J.S., Gilroy, L.A., Blake, R., 2003. Perceptual consequences of centre-surround antagonism in visual motion processing. Nature 424, 312–315.

Tadin, D., Park, W.J., Dieter, K.C., Melnick, M.D., Lappin, J.S., Blake, R., 2019. Spatial suppression promotes rapid figure-ground segmentation of moving objects. Nature Communications.

Tadin, D., Silvanto, J., Pascual-Leone, A., Battelli, L., 2011. Improved Motion Perception and Impaired Spatial Suppression following Disruption of Cortical Area MT/V5. J Neurosci 31, 1279–1283.

Takada, N., Pi, H.J., Sousa, V.H., Waters, J., Fishell, G., Kepecs, A., Osten, P., 2014. A developmental cell-type switch in cortical interneurons leads to a selective defect in cortical oscillations. Nature Communications.

Tan, H.R.M., Gross, J., Uhlhaas, P.J., 2016. MEG sensor and source measures of visually induced gamma-band oscillations are highly reliable. Neuroimage 137, 34–44.

Taulu, S., Hari, R., 2009. Removal of magnetoencephalographic artifacts with temporal signal-space separation: demonstration with single-trial auditory-evoked responses. Hum Brain Mapp 30, 1524–1534.

Troche, S.J., Thomas, P., Tadin, D., Rammsayer, T.H., 2018. On the relationship between spatial suppression, speed of information processing, and psychometric intelligence. Intelligence 67, 11–18.

van Pelt, S., Boomsma, D.I., Fries, P., 2012. Magnetoencephalography in Twins Reveals a Strong Genetic Determination of the Peak Frequency of Visually Induced Gamma-Band Synchronization. J Neurosci 32, 3388–3392.

van Pelt, S., Fries, P., 2013. Visual stimulus eccentricity affects human gamma peak frequency. Neuroimage 78, 439–447.

van Pelt, S., Shumskaya, E., Fries, P., 2018. Cortical volume and sex influence visual gamma. Neuroimage 178, 702–712.

Whithain, E.M., Pope, K.J., Fitzgibbon, S.P., Lewis, T., Clark, C.R., Loveless, S., Broberg, M., Wallace, A., DeLosAngeles, D., Lillie, P., Hardy, A., Fronsko, R., Pulbrook, A., Willoughby, J.O., 2007. Scalp electrical recording during paralysis: Quantitative evidence that EEG frequencies above 20 Hz are contaminated by EMG. Clinical Neurophysiology 118, 1877–1888.

Whitham, E.M., Lewis, T., Pope, K.J., Fitzgibbon, S.P., Clark, C.R., Loveless, S., DeLosAngeles, D., Wallace, A.K., Broberg, M., Willoughby, J.O., 2008. Thinking activates EMG in scalp electrical recordings. Clinical Neurophysiology 119, 1166–1175.

Zhu, M.C., Rozell, C.J., 2013. Visual Nonclassical Receptive Field Effects Emerge from Sparse Coding in a Dynamical System. Plos Computational Biology.

